# Mobile immune signals potentiate salicylic acid-mediated plant immunity via WRKY38/62 transcription factors

**DOI:** 10.1101/2025.04.17.649115

**Authors:** Robert O. Mason, Heather Grey, Steven H. Spoel

## Abstract

Systemic acquired resistance (SAR) is a broad-spectrum plant immune response that provides protection against various pathogens. Activation of SAR requires mobile immune signals as well as the indispensable immune hormone salicylic acid (SA). Nonetheless, it remains unknown how mobile signals integrate with the SA signal to produce functional SAR responses. Here, we demonstrate that the mobile signals, azelaic acid (AzA) and N-hydroxy-pipecolic acid (NHP), respectively dampened and potentiated SA-induced transcriptional reprogramming of thousands of genes. Indeed, NHP enhanced stability of the SA receptor protein NPR1, and unlike AzA, it dramatically increased the effectiveness of SA-induced immunity against bacterial infection by 10-fold. Analysis of NHP-primed, SA-responsive gene promoters indicated that WRKY transcription factors play an important role in integrating these two immune signals. While responsiveness to SA remained largely unaffected by mutation of *WRKY38* and *WRKY62*, it abolished NHP-mediated potentiation of SA-induced gene expression and immunity. Collectively, our findings reveal mobile signals potentiate SA-mediated plant immunity via WRKY38/62 transcription factors.

**Teaser:** Activation of broad-spectrum immunity in plants requires mutual potentiation between multiple distinct immune signals.

## Introduction

Like all eukaryotes, plants are constantly exposed to a variety of pathogenic threats and have evolved sophisticated immune responses to fend off invaders. Plant immune responses are characterized by dramatic reprogramming of the transcriptome to prioritize immune responses over other cellular functions. This process is largely regulated by the immune hormone, salicylic acid (SA), which accumulates upon pathogen infection (*1-3*). SA induces not only local resistance, but also systemic acquired resistance (SAR), a long-lasting immune response with broad-spectrum effectiveness (*4*).

SA is perceived by various different receptor proteins that regulate distinct cellular processes (*5*), including by members of the non-expressor of pathogenesis-related genes (NPR) family (*6-10*). Perception of SA by the NPR1 protein is arguably the most important for establishment of immunity. NPR1 forms nuclear condensates where it acts as a coactivator of SA-responsive genes by establishing a transcriptional activation complex with TGA transcription factors and chromatin modifiers (*11-13*). NPR1 is tightly regulated by a series of post-translational modifications that modulate its activity and stability (*5*). In particular, NPR1 is sequentially modified in a relay by three different types of ubiquitin ligases. Nuclear ubiquitination by a Cullin-RING Ligase 3 (CRL3) promotes NPR1 coactivator activity and chromatin association, while subsequent ubiquitin chain extension by ubiquitin conjugation factor E4 (UBE4) deactivates NPR1 and recruits the proteasome (*14, 15*). Finally, modification by the proteasome-associated HECT-type ubiquitin protein ligase 3 (UPL3) and UPL4 is required for processive degradation of NPR1 (*16, 17*). Conversely, these ubiquitin ligase activities are countered by dedicated deubiquitinases, allowing dynamic ubiquitination of NPR1 to fine-tune its transcription coactivator activity (*15*).

In addition to SA, establishment of SAR requires mobile signals that travel through the vasculature from the site of infection to systemic tissues, where they promote immune priming through mostly uncharacterized mechanisms (*4, 18, 19*). These mobile signals include free radicals (*20*), glycerol-3-phosphate (*21*), dehydroabietinal (*22*), azelaic acid (AzA)(*23*) and N-hydroxy-pipecolic acid (NHP)(*24*). Both AzA and NHP have been suggested to be linked to SA signaling. On the one hand, lipid-derived Aza accumulates upon infection, is transported systemically throughout the plant by Azelaic Acid Induced 1 (AZI1) that resembles a lipid transfer protein, and primes the accumulation of SA (*19, 23, 25*). On the other, NHP is synthesized from lysine through the intermediary pipecolic acid (Pip) by the enzymes AGD2-like defense response protein (ALD1), SAR deficient 4 (SARD4) and flavin-dependent monooxygenase 1 (FMO1)(*24, 26, 27*). Exogenous treatment of lower leaves with NHP increased pathogen-mediated production of SA, NHP and camalexins, and is sufficient to confer resistance to infection in upper leaves (*28*). Moreover, exogenous NHP triggers transcriptional changes that are reminiscent to those observed during SAR. This requires SA biosynthesis, NPR1 and TGA transcription factors (*28, 29*), suggesting that NHP cooperates with the SA signaling pathway to induce SAR. Nonetheless, it remains unknown how both Aza and NHP promote SA-dependent SAR.

Here, we demonstrate that while Aza dampens SA signaling, NHP strongly potentiates and reprograms SA-induced, genome-wide transcriptional responses. Consequently, NHP increased the effectiveness of SA against bacterial infection by 10-fold. We show that NHP potentiates SA-induced immunity by stabilizing NPR1 protein and identified two WRKY transcription factors that are indispensable for the priming effects of NHP.

## Results

### AzA dampens SA-induced transcriptional immune reprogramming

To investigate if mobile signals interact with the SA signal, we examined if mobile signals modified SA-induced immune gene expression. Unlike application of SA, treatment of wild-type *Arabidopsis* plants with AzA did not elicit expression of SA-responsive immune genes (Fig. 1A). However, pre-treatment with AzA strongly enhanced the SA-induced expression of *PR1*, but not *WRKY38 and WRKY62* (Fig. 1A).

**Fig. 1.**
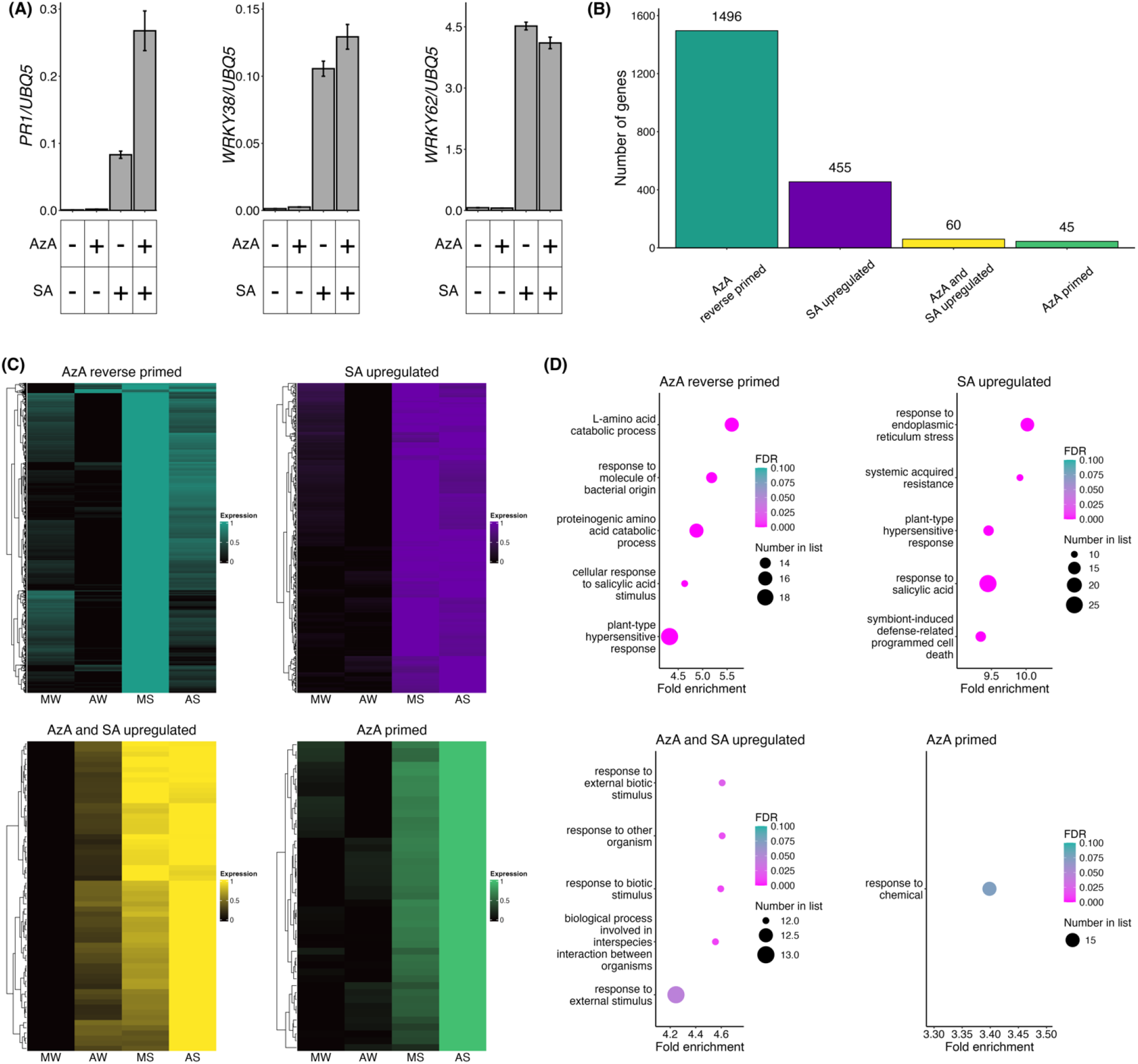
AzA dampens SA-induced transcriptional reprogramming. (**A**) AzA selectively primes expression of the SA-induced reporter genes *PR1, WRKY38* and *WRKY62*. Reporter genes were normalised against constitutively expressed *UBQ5*. (**B**) RNA-seq clusters of AzA-modified, SA-induced genes. Clusters were identified and assigned by hierarchical clustering using the hclust R package. (**C**) Heatmaps of clustered genes with expression patterns across the four treatment conditions (MW: mock + water, AW: AzA + water, MS: mock + SA, AS: AzA + SA). (**D**) Top 5 most enriched GO terms in each cluster (p ≤ 0.05, FDR ≤ 0.1). For all experiments, 14-day-old wild-type seedlings were treated by immersion in 5 mM MES (pH 5.6) with or without 1 mM AzA for 18 hours, followed by treatment with water or 0.5 mM SA for 6 hours. Total RNA was extracted and the expression of SA-responsive genes measured by qPCR or RNA-seq.

To further characterize if AzA alters SA-mediated transcriptional reprogramming, we performed RNA sequencing on plants sequentially treated with or without AzA and then SA. AzA alone specifically altered the expression of only 647 genes (Table S1A). However, when sequentially applied with SA, AzA strikingly modified 75% of SA-responsive genes, suggesting its major role is to modify SA-mediated transcriptional reprogramming (Fig 1B, Table S1B). SA-regulated genes were clustered according to expression changes triggered by pre-treatment with AzA. This broadly identified four expression categories for genes that were induced by SA (Fig. 1B, C). The largest of these categories contained 1,496 differentially expressed genes (DEGs) that exhibited reduced SA-induced expression in presence of AzA (‘AzA reverse primed’; Fig. 1B, C). GO analyses of these genes demonstrated that AzA primarily suppressed SA-responsive genes involved in responses to bacterial pathogens and in hypersensitive programmed cell death responses (Fig 1D). By contrast, only a handful of genes characterised by their response to various stimuli, were either upregulated by both SA and AzA (60 DEGs), or displayed enhanced SA-induced expression after AzA treatment (45 DEGs)(Fig 1B-1D). These results demonstrate that AzA largely counters the SA-mediated upregulation of immune genes, potentially to avoid programmed cell death responses that during SAR are undesirable in systemic tissues.

Conversely, the effect of AzA was more varied on genes that were repressed by SA. While AzA countered the SA-mediated downregulation of 406 genes, it also facilitated the repression of 672 genes (fig S1A, B and Table S1B). This outnumbered the 391 genes whose SA-mediated downregulation was primed or reverse primed by AzA (184 and 207, respectively). Notably, GO analyses showed that AzA opposed SA in the down regulation of cellular household functions, including photosynthesis and metabolism (fig S1C). Taken together, our findings suggest that AzA primes plants primarily by dampening the effect of SA on a variety of processes, including its role as a cell death agonist, while also limiting its role in prioritising immunity over other cellular activities.

### NHP potentiates SA-induced transcriptional immune reprogramming

Next, we examined if like AzA, the major mobile immune signal NHP is capable of modifying SA-mediated gene expression. While application of NHP alone did not induce SA-responsive marker genes, sequential treatment with NHP and SA dramatically enhanced SA-induced expression of all three marker genes examined (Fig. 2A), suggesting NHP may potentiate SA-induced transcriptional reprogramming.

**Fig. 2.**
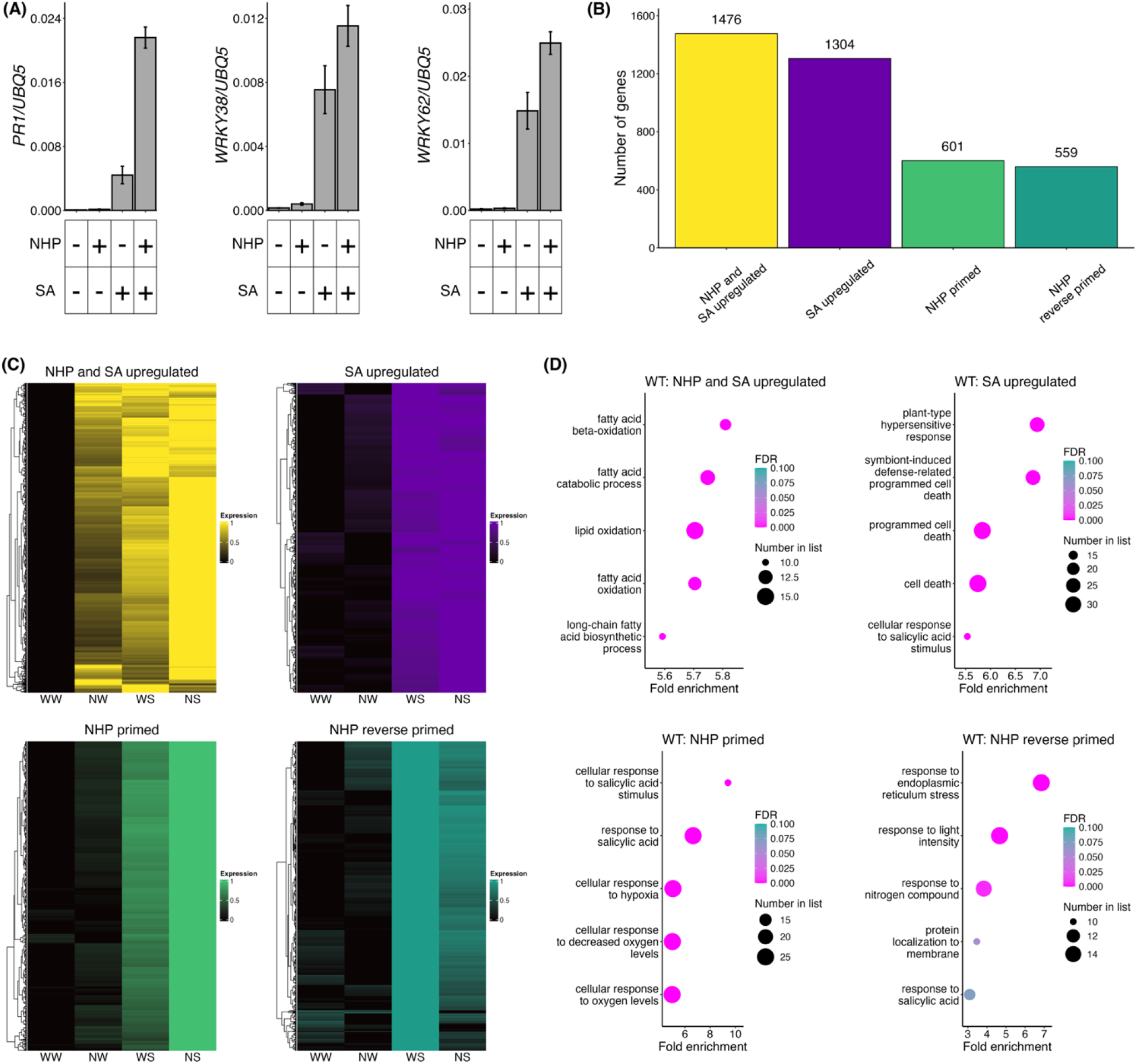
NHP primes SA-induced transcriptional reprogramming. (**A**) NHP potentiates expression of the SA-induced reporter genes *PR1, WRKY38* and *WRKY62*. Reporter genes were normalised against constitutively expressed *UBQ5*. (**B**) RNA-seq clusters of NHP-modified, SA-induced genes. Clusters were identified and assigned by hierarchical clustering using the hclust R package. (**C**) Heatmaps of clustered genes with expression patterns across the four treatment conditions (WW: water + water, NW: NHP + water, WS: water + SA, NS: NHP + SA). (**D**) Top 5 most enriched GO terms in each cluster (p ≤ 0.05, FDR ≤ 0.1). For all experiments, 14-day-old wild-type seedlings were treated by immersion in water or 1 mM NHP for 18 hours, followed by treatment with water or 0.5 mM SA for 6 hours. Total RNA was extracted and the expression of SA-responsive genes measured by qPCR or RNA-seq.

To investigate this further, we examined the transcriptomes of plants sequentially treated with or without NHP and then SA. Only 844 genes showed expression changes specifically to NHP alone (Table S2A), compared to 2,762 SA-induced genes and 2,349 SA-repressed genes that were modified by NHP (Fig. 2B, fig S2A, Table S2B). Indeed, 67% of SA-mediated genes that were upregulated and 87% of downregulated genes were modulated by NHP. To understand how NHP modifies SA signalling, we clustered SA-regulated genes according to expression changes caused by pre-treatment with NHP. This uncovered 1,476 genes that were synergistically induced by NHP and SA, and another 601 genes whose SA-induced expression was primed by NHP (Fig 2B, 2C). These NHP-adjusted genes formed typical GO categories associated with responsiveness to SA, but excluded terms associated with SA-induced programmed cell death (Fig 2D). Conversely, 1,290 genes were repressed in a synergistic manner by NHP and SA, along with a modest 128 genes whose SA-mediated downregulation was potentiated in presence of NHP (Fig S2A, S2B). Nevertheless, NHP played more subtle roles beyond only potentiating SA-responsive genes. It also ‘reverse primed’ many SA-responsive genes, so that these genes exhibited reduced SA-mediated expression changes after NHP treatment (Fig 2B, 2C). These genes primarily displayed GO terms associated with translation and ribosomal RNA processing (Fig 2D). Moreover, a substantial number of 678 genes were regulated in opposite manner by NHP and SA, thereby dampening their SA-mediated downregulation (fig S2). Collectively, these data strongly imply that potentiation and priming to fine-tune SA-responsive gene expression is the major role of NHP.

### NHP but not AzA is sufficient to amplify SA-induced disease resistance

To determine if AzA- and NHP-mediated changes in SA-responsive gene expression correspond to enhanced immune responses, we performed disease assays with the bacterial leaf pathogen *Pseudomonas syringae* pv. *maculicola* (*Psm*) ES4326. Foliar treatment with either AzA or with the NHP precursor, pipecolic acid (Pip) that also potentiates SA responses (fig S3), did not induce resistance against *Psm* ES4326 (Fig 3A, 3B). Similarly, treatment with a low concentration of SA (0.05 mM) also did not induce resistance, while a 10-fold higher concentration of SA (0.5 mM) induced strong protection against infection. Strikingly, while AzA was unable to enhance the effectiveness of low concentrations of SA (Fig 3A), sequential treatment with Pip and 0.05 mM SA strongly limited pathogen growth to a similar level of protection as application of a ten-fold higher concentration of SA alone (Fig 3B). These data demonstrate that only NHP-mediated priming of SA-induced transcriptional reprogramming renders previously inert concentrations of SA sufficient to induce disease resistance against *Psm* ES4326.

**Fig. 3.**
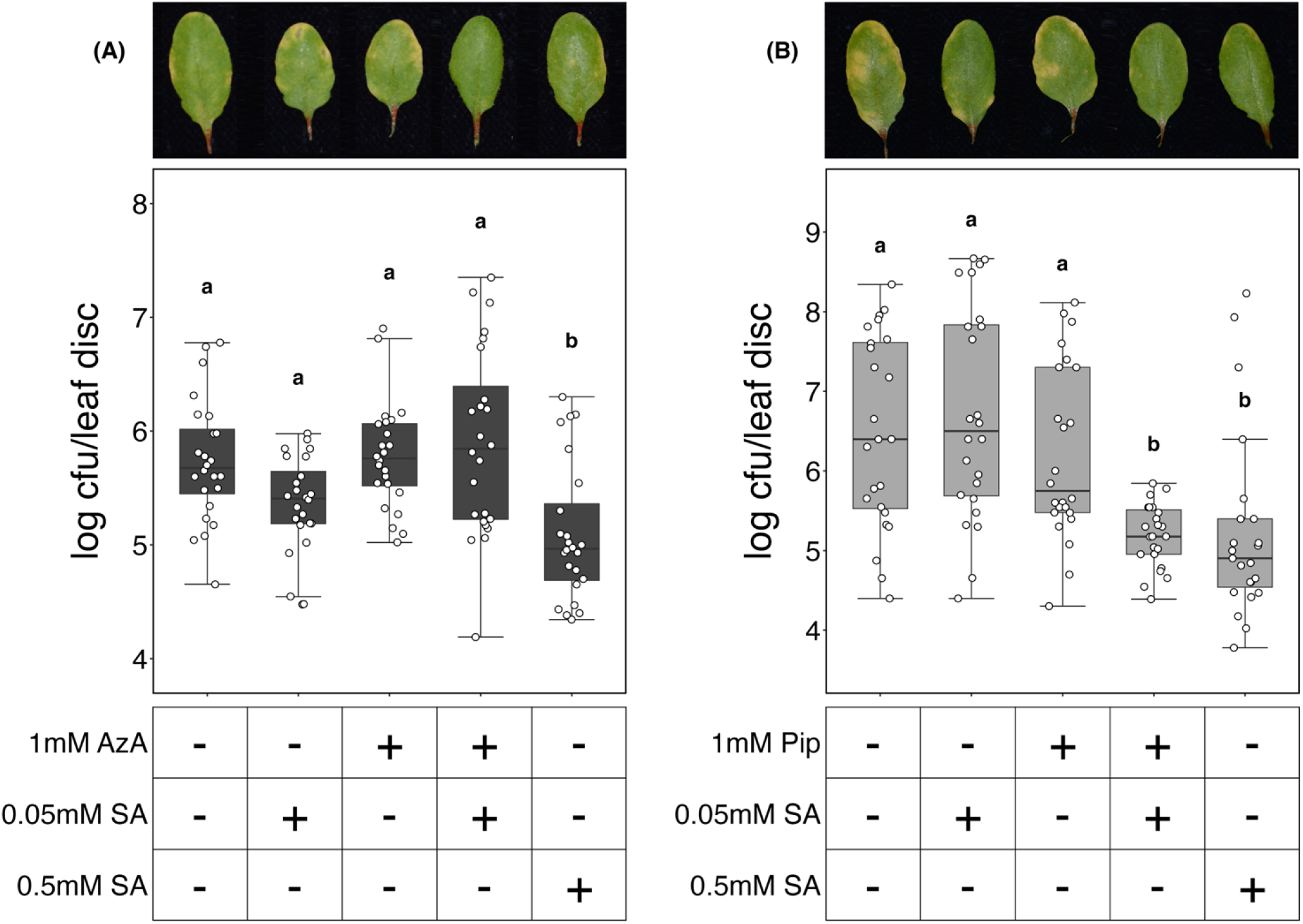
NHP but not AzA primes SA-induced immunity against *Psm* ES4326. (**A**) Pre-treatment with AzA does not prime SA-induced immunity against *Psm* ES4326. (**B**) Pip-mediated priming renders previously inert concentrations of SA sufficient to induce disease resistance against *Psm* ES4326. Twenty-eight day-old wild-type plants were treated 48 hours before infection by spraying with 5 mM MES (pH 5.6) supplemented with or without 1 mM AzA or with water supplemented with or without 1 mM Pip, followed by treatment with water, 0.05 mM SA or 0.5 mM SA, 24 hours before infection. Leaves were then infiltrated with 5 x 10^6^ colony forming units (cfu)/ml *Psm* ES4326. Leaf discs were analysed for bacterial growth 3 days post infection. Error bars represent interquartile range x1.5, while letters denote statistically significant differences between samples (Tukey ANOVA; α = 0.05, n = 24). Representative photos of leaves are displayed above the respective boxplots.

### NHP primes NPR1 protein levels by reducing its turnover

Because SA-induced gene expression reprogramming is dependent on the SA receptor and transcription coactivator NPR1, we examined if NHP affects steady-state levels of NPR1 protein. *35S:NPR1-GFP* (in *npr1-1*) plants were treated with NHP, SA or a sequential combination of both. Similar to SA treatment, application of NHP induced accumulation of NPR1-GFP protein (Fig 4A), independent of mRNA expression levels (Fig 4B). Sequential treatment with both signals did not further enhance NPR1-GFP accumulation beyond treatment with SA alone. NPR1 is subject to proteasome-mediated degradation, which regulates both its steady-state and SA-activated protein levels (*14, 15, 17*). Therefore, it is plausible that NHP enhanced NPR1 protein levels by increasing its stability. To test this, plants were treated with the protein synthesis inhibitor, cycloheximide (CHX), and the stability of NPR1-GFP monitored through time. Compared to mock treatment, NPR1-GFP protein was degraded at a slower rate in presence of NHP (Fig 4C), indicating NHP stabilised NPR1-GFP. In presence of SA, turnover of NPR1 is coupled to its ability to activate target genes (*14, 15*). In accordance, NHP did not stabilise SA-activated NPR1-GFP protein (Fig 4D). These findings suggest that NHP potentiates SA-mediated transcriptional reprogramming by elevating the levels of NPR1 protein prior to full activation of SA signalling.

**Fig. 4.**
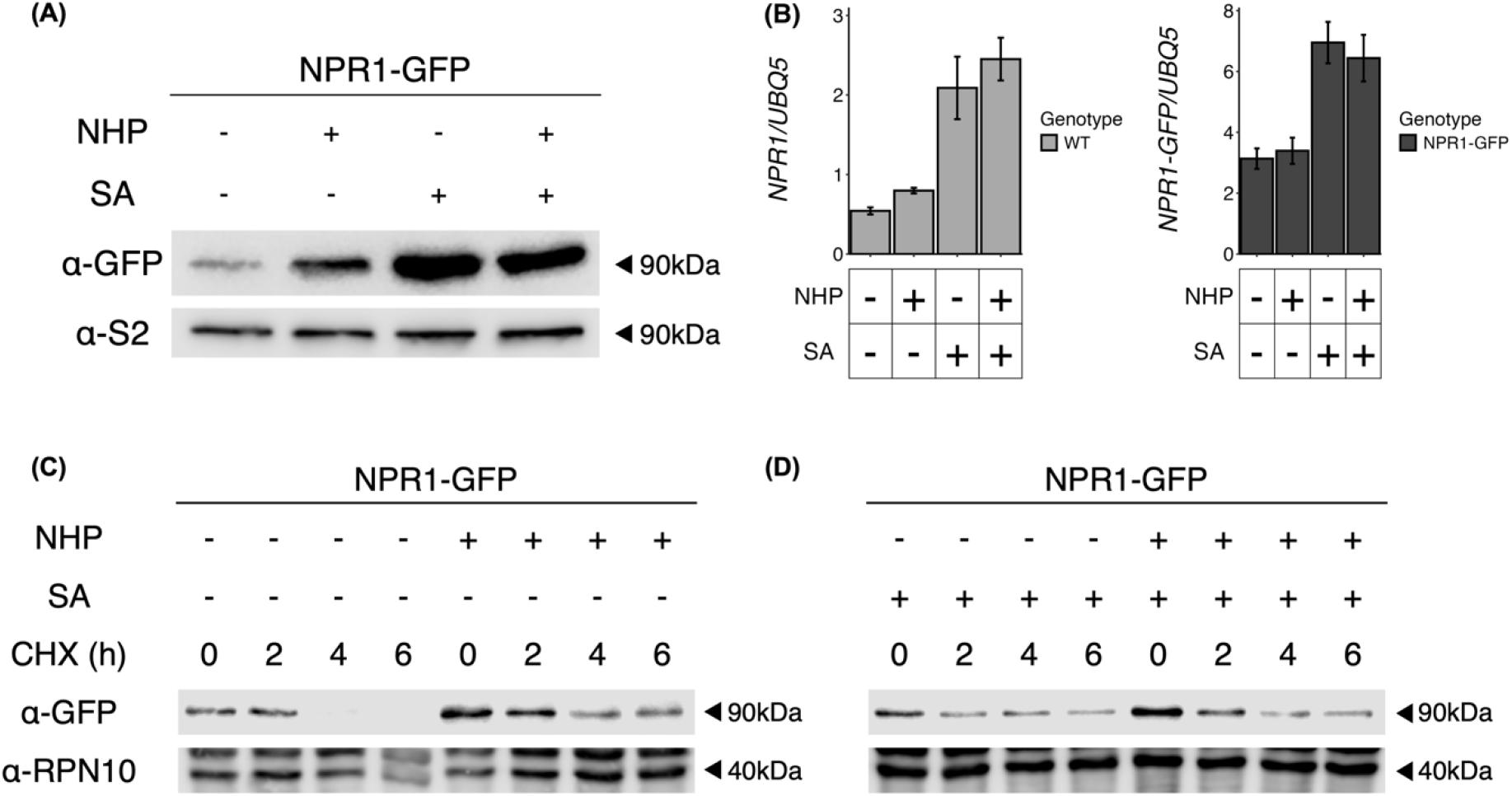
NHP stabilises and elevates NPR1 protein levels. (**A**) NHP treatment induces NPR1 accumulation but does not enhance SA-mediated NPR1 accumulation. Fourteen-day-old *35S::NPR1-GFP* (*npr1*) seedlings were treated by immersion in water or 1 mM NHP for 18 hours, followed by treatment with water or 0.5 mM SA for 6 hours. Total protein was extracted and separated by electrophoresis, NPR1 was visualised with an anti-GFP antibody and inputs visualised with anti-S2. (**B**) NHP treatment does not induce or prime expression of the endogenous *NPR1* gene in wild type plants or of the *NPR1-GFP* transgene in *35S:NPR1-GFP* (*npr1*) plants. Fourteen-day-old seedlings were treated by immersion in water or 1 mM NHP for 18 hours, followed by treatment with water or 0.5 mM SA for 6 hours. Total RNA was extracted, the expression of *NPR1* and *NPR1-GFP* genes measured by qPCR and normalised against constitutively expressed *UBQ5*. (**C**) NHP treatment stabilises NPR1 protein levels, but (**D**) NHP does not enhance SA-mediated NPR1 stability. *35S:NPR1-GFP* seedlings were treated as described in (A), but 2h after addition of SA, 100 µM CHX was added. Samples were taken immediately after adding CHX and thereafter at 2h, 4h and 6h timepoints. Total protein was extracted and analysed with anti-GFP and anti-RPN10 antibodies.

Overexpression of *NPR1* has been reported to elevate the expression of its target genes (*30*). To test if elevated levels of NPR1 protein are sufficient to substitute for NHP-mediated potentiation of SA-responsive gene expression, we employed mutants of ubiquitin ligases that target NPR1 for degradation. Mutants of either Cullin 3 (*CUL3*) E3 ligase or both its substrate adaptors, NPR3 and NPR4, as well as the E4 ligase UBE4, all accumulate elevated levels of NPR1 protein (*6, 14, 15*). Nonetheless, we found that NHP continued to prime expression of the SA-induced *PR1* gene in all three mutants (Fig 5). Thus, while NHP promotes NPR1 protein accumulation, this is in itself insufficient to potentiate SA-responsive gene expression.

**Fig. 5.**
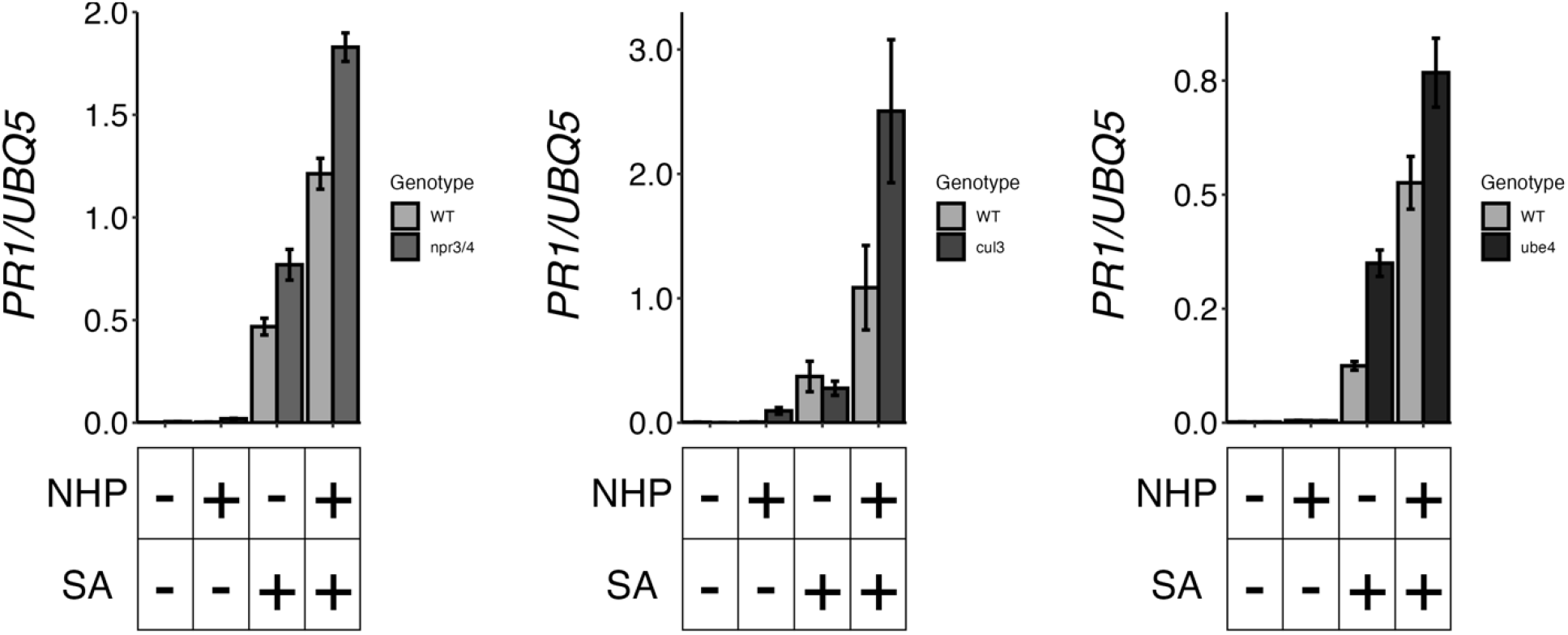
Elevated levels of NPR1 protein are insufficient to substitute for NHP-mediated priming of SA-responsive gene expression. Similar to wild-type (WT), application of NHP potentiated SA-induced *PR1* gene expression in *npr3/4, cul3a/b* (abbreviated as *cul3*), and *ube4* mutants that all have elevated levels of NPR1 protein. Fourteen-day-old seedlings were treated by immersion in water or 1 mM NHP for 18 hours, followed by treatment with water or 0.5 mM SA for 6 hours. Total RNA was extracted and the expression of SA-responsive *PR1* measured by qPCR and normalised against constitutively expressed *UBQ5*.

### WRKY38 and WRKY62 are master regulators of NHP-mediated potentiation of SA signalling

Because NPR1 protein levels alone do not explain how NHP potentiates SA signalling, we analysed transcription factors binding sites in the promoters of NHP-potentiated genes. Using Chi squared tests, we identified transcription factor interactions that were overrepresented compared to all DEGs. Using published ChIP-seq and DAP-seq datasets (*31*), we found that WRKY transcription factors were highly abundant, with 17 of the top 20 motifs belonging to this family (Table S3). In particular this revealed potential roles for WRKY18, WRKY54 and WRKY70, all of which have been reported to play integral roles in modulating the immune transcriptome.

WRKY18 acts as an NPR1-induced auxiliary transcription factor that acts to amplify SA-responsive gene expression and defence (*14, 32*). To test if NHP utilises WRKY18 to amplify SA signalling, we treated *wrky18* mutants sequentially with NHP and SA. However, NHP was still able to potentiate the SA-induced expression of *PR1* (Fig 6), indicating it does not play a major role in priming. We then considered WRKY70, which together with WRKY54 act as positive regulators of SA-mediated gene expression and immunity (*32*). Moreover, WRK54/70 were recently found to be induced early after NHP treatment and WRKY70 was required for NHP-induced disease resistance (*33*). Thus, we examined if *wrky54/70* mutants failed to exhibit NHP-mediated potentiation of SA-responsive gene expression. While SA-induced *PR1* gene expression was dramatically reduced in *wrky54/70* mutants, co-treatment with NHP still potentiated this SA response (Fig 6). This suggests that although WRKY54/70 are important for both SA- and NHP-dependent responses, they are not the main contributors to NHP’s ability to potentiate SA signalling. Lastly, we examined the involvement of WRKY38 and WRKY62. Curiously, although SA-induced resistance to *Psm* ES4326 was comparable between *wrky38/62* double mutants and wild-type (fig S4), these mutants were previously reported to lack activation of true SAR induced by an avirulent pathogen (*14*). It is therefore plausible that WRKY38/62 specifically regulate defence priming. Indeed, whereas *wrky38/62* mutants were not impaired in SA-induced *PR1* gene expression, NHP-mediated potentiation of this response was completely abolished (Fig 6). Thus, WRKY38/62 may be master regulators of NHP-mediated potentiation of SA signalling.

**Fig. 6.**
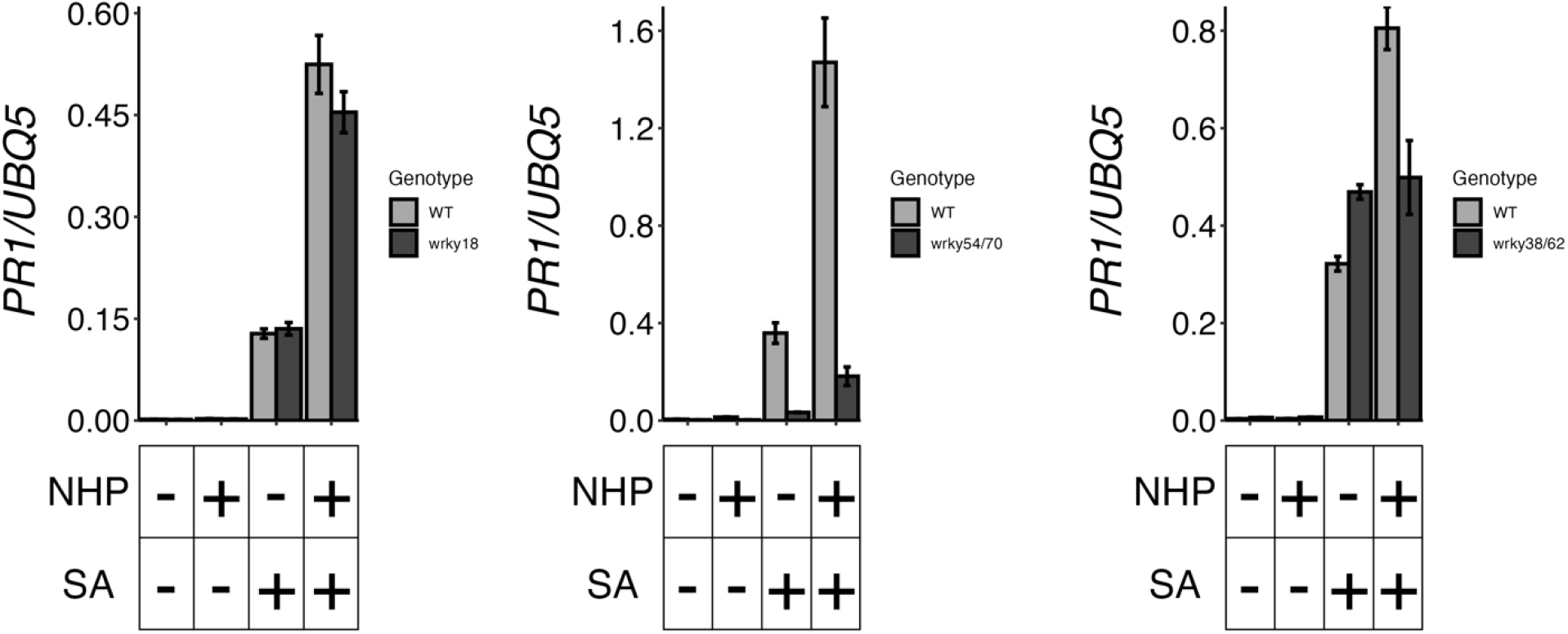
NHP-mediated potentiation of SA-responsive *PR1* gene expression is differentially regulated by WRKY transcription factors. NHP-mediated potentiation of SA-induced *PR1* gene expression was largely intact in *wrky18* and *wrky54/70* mutants, but is completely abolished in *wrky38/62* mutants. Fourteen-day-old seedlings were treated by immersion in water or 1 mM NHP for 18 hours, followed by treatment with water or 0.5 mM SA for 6 hours. Total RNA was extracted and the expression of SA-responsive *PR1* measured by qPCR and normalised against constitutively expressed *UBQ5*.

### Potentiation of SA-induced transcriptional reprogramming and immunity by NHP requires WRKY38 and WRKY62

To determine if the loss of NHP-mediated priming in *wrky38/62* mutants was reflected across the entire SA-responsive transcriptome, RNA-seq was performed to compare these mutants to wild type. The results paint a complex but clear broad picture. First, in *wrky38/62* mutants, over 70% of SA-induced genes modulated by NHP in the wild type, now became only responsive to SA or were rendered completely unresponsive to either NHP or SA (Fig 7A). Similarly, in absence of functional *WRKY38/62*, NHP also largely failed to modulate the expression of genes downregulated by SA (Fig 7B). Thus, WRKY38/62 are required for global potentiation of the SA-responsive transcriptome. Second, a smaller group of SA-induced genes experienced misregulation by NHP in *wrky38/62* mutants. These genes typically switched between the NHP primed, reverse primed and synergistic regulatory clusters (Fig 7A). A similar phenomenon was observed for genes down regulated by SA (Fig 7B). Together, these findings demonstrate that WRKY38/62 are master regulators of NHP-mediated potentiation of SA-induced transcriptional reprogramming.

**Fig. 7.**
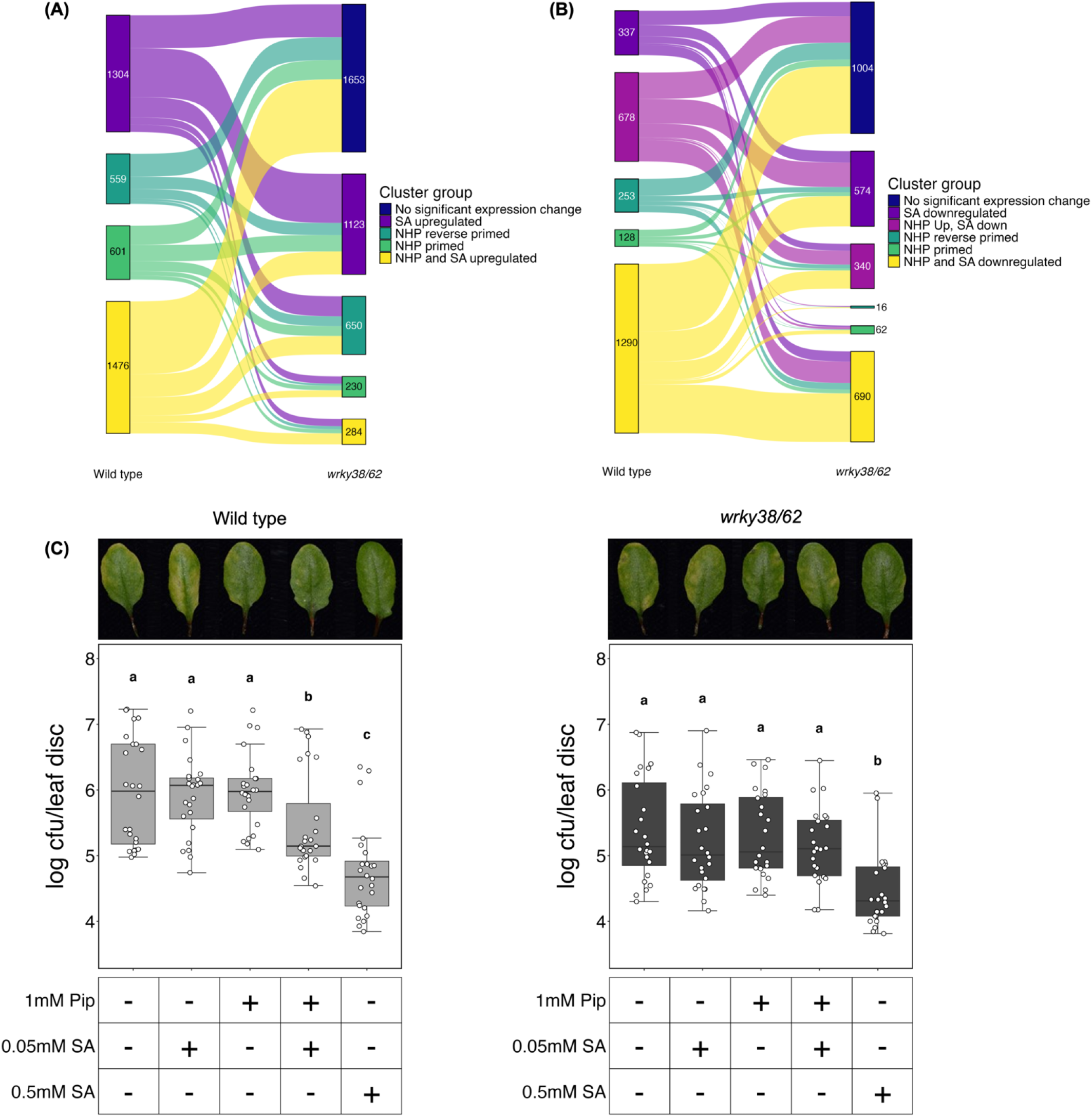
NHP-mediated priming of SA-induced transcriptional reprogramming and immunity requires WRKY38 and WRKY62. (**A**) Sankey flow diagram of the behaviour of different NHP-modified clusters of SA-upregulated genes in WT and *wrky38/62* mutant plants. Note that the “SA upregulated” clusters refer to genes that were induced by SA but their expression was not influenced by NHP, while “No significant expression change” indicates genes that failed to display differential expression in *wrky38/62* mutants. (**B**) Sankey flow diagram of the behaviour of different NHP-modified clusters of SA-downregulated genes in WT and *wrky38/62* mutant plants. Clusters are as in panel (A) but for genes downregulated by SA. For data in (A) and (B) 14-day-old seedlings were treated by immersion in water or 1 mM NHP for 18 hours, followed by treatment with water or 0.5 mM SA for 6 hours. Total RNA was extracted and the expression of SA-responsive genes measured by RNA-seq. (**C**) Pip-mediated priming of disease resistance against *Psm* ES4326 is lost in *wrky38/62* mutants. Twenty-eight day-old wild-type or *wrky38/wrky62* plants were treated by spraying with water or 1 mM Pip, 48 hours before infection, followed by treatment with water, 0.05 mM SA or 0.5 mM SA, 24 hours before infection. Leaves were then infiltrated with 5 x 10^6^ colony forming units (cfu)/ml *Psm* ES4326. Leaf discs were analysed for bacterial growth two days post infection. Error bars represent interquartile range x 1.5 limits, while letters denote statistically significant differences between samples (Tukey ANOVA; α = 0.05, n = 24). Representative photos of leaves are displayed above their respective boxplots.

Because of the dramatic changes observed in the NHP- and SA-responsive transcriptome of *wrky38/62* mutants, we investigated if these WRKYs conferred NHP-elicited amplification of SA-induced immunity. As described earlier, foliar treatment with the NHP precursor, Pip, or with low concentrations of SA (0.05 mM) failed to induce resistance against *Psm* ES4326 in both wild-type and mutant *wrky38/62* plants (Fig 7C). By contrast, treatment with a ten-fold higher concentration of SA (0.5 mM) protected both the wild type and mutant from infection. Sequential treatment with Pip and 0.05 mM SA, however, limited pathogen growth only in the wild type and not in *wrky38/62* mutants (Fig 7C). Taken together, our results demonstrate that NHP utilises WRKY38/62 for potentiation of the SA-responsive transcriptome to establish SAR.

## Discussion

NHP and AzA have been identified as key mobile signals in plant immunity, but how they interact with other immune signals is not well understood. Here, we show that AzA modulates SA-responsive gene expression, but is not sufficient to alter immune phenotypes. By contrast, we demonstrate that NHP primes SA-induced transcriptional reprogramming and immunity by employing the WRKY38 and WRKY62 transcription factors.

Compared to NHP, less is known about how AzA contributes to immunity and what its relationship is with SA. AzA is produced in response to pathogen infection, accumulating in the phloem and travelling to systemic tissues (*23*). However, reports on its role in immunity are inconsistent. For example, it has been reported that AzA accumulates in response to infection in tobacco, but exogenous application alone was insufficient to induce resistance (*34*). Conversely, it has been shown that application of AzA to *Arabidopsis* leaves or roots enhances resistance to *P. syringae* in aerial parts, but without apparent transport of AzA (*23, 25*). These inconsistencies may point towards AzA being only a single component of a wider signature of interacting immune signals. In this study we considered that after infection, AzA and SA both accumulate in systemic tissues (*1, 2, 23*), and therefore may cross talk. Indeed, we found that AzA primarily dampened SA-induced transcriptional reprogramming related to hypersensitive cell death, while also limiting SA’s ability to suppress cellular ‘household’ activities (Fig 1, fig S1). Hypersensitive cell death is a vital response for local tissues to halt pathogen infection, but is undesirable in systemic tissues where long-lasting SAR is established. Moreover, the energy requirements for establishing SAR are probably much lower than those of immune responses in local tissues, so prioritisation of immunity over other cellular functions may be less dramatic in systemic tissues. So, while AzA alone was insufficient to enhance SA-induced immunity against infection (Fig 3A), its ability to tailor the SA-responsive transcriptome in systemic cells is likely a key facet of SAR.

Of all identified mobile immune signals to date, NHP shows the strongest relationship with SA. Upon infection, the transcription factors SARD1 and CBP60g activate both NHP and SA biosynthesis in infected cells (*35*). NHP is transported from the site of infection to systemic tissues, where it induces the biosynthesis of SA as well as itself, enabling these two immune signals to promote development of SAR (*28*). NHP also primes signalling downstream of SA biosynthesis, as Pip or NHP treatment both enhanced SA-induced *PR1* gene expression (*28, 36*). After corroborating these findings here with additional SA-responsive marker genes (Fig 2A, fig S3), we explored transcriptome-wide interplay between NHP and SA. We discovered that NHP’s main role is to modify the SA-responsive transcriptome, with over three-quarters of SA-responsive genes exhibiting markedly altered expression in presence of NHP (Fig 2, fig S2). The majority of these genes were either mutually regulated by both NHP and SA or their SA-responsiveness was primed by NHP. Consequently, NHP-elicited potentiation of the SA-responsive transcriptome was associated with a dramatic 10-fold increase in effectiveness of SA-induced immunity against bacterial infection (Fig 3). Our findings therefore explain how low levels of SA observed in systemic tissues can launch a highly effective immune response.

In addition to priming SA-responsive genes for enhanced induction or repression, NHP also reverse primed a substantial set of genes (*i*.*e*. they became less responsive to SA) (Fig 2, fig S2). GO analyses suggested that these genes are involved in light and nutrient signalling or translation (Figs 2D and S2C). Thus, in a similar manner to AzA’s dampening effect on SA signalling, NHP also aids in tailoring the immune transcriptome specifically for systemic tissues. NHP may further determine the behaviour of systemic tissues by promoting the stabilisation of NPR1 protein and enhancing its accumulation (Fig 4)(*37*). Notably, cell death triggered by intracellular immune receptors in infected local tissues is curtailed by accumulation of NPR1 protein in neighbouring cells (*6, 11*). Here, NPR1 localises to cytoplasmic SA-induced condensates (known as cSINC) where it is thought to target cell death-inducing immune receptors and their downstream signalling components for degradation, thereby promoting cell survival (*11, 38*). In combination with AzA’s ability to suppress cell death-related genes, NHP-induced NPR1 protein levels may therefore ensure survival of systemic tissues. This may be a particularly important attribute to ensure that NHP’s ability to induce SA synthesis (*28, 29*) and dramatically enhance SA signalling (Figs 2 and 3), does not escalate into undesirable hypersensitive cell death in systemic tissues.

NPR1 is required for both SA- and NHP-mediated immunity, but is likely not a direct receptor of NHP (*28, 29, 37, 39*). So does NHP prime SA-responsive gene expression simply by increasing NPR1 protein levels? In *Arabidopsis*, engineering increased expression levels of NPR1 protein leads to stronger SA-induced *PR* gene expression and resistance to pathogens apparently without major growth trade-offs (*30*). Because overexpression of NPR1 resembles a priming response, we investigated if elevating NPR1 protein levels by mutating E3 ligases that target NPR1 for degradation (*6, 14, 15*), can substitute for NHP. Surprisingly, NHP still induced strong priming responses in *npr3/4, cul3* and *ube4* mutants (Fig 5), indicating that enhanced NPR1 protein levels alone are insufficient to confer NHP-mediated priming of SA responses. To uncover how NHP primes SA-responsive genes, we turned our attention to promoter motifs in primed genes. To our initial disappointment, we found that WRKY transcription factor-binding motifs were highly overrepresented in primed gene promoters (Table S3). WRKY transcription factors are well known to regulate SA-responsive gene expression downstream of NPR1 (*32*), so their identification was less than surprising. Indeed, WRKY18, which acts as a key auxiliary transcription factor to amplify SA- and NPR1-dependent gene expression (*32*), was not required for NHP-elicited priming of SA-responsive gene expression (Fig 6). However, WRKYs have recently also been linked with the regulation of NHP signalling. In particular, WRKY54 and WRKY70 were found to be required for NHP-induced gene expression and SAR (*33*). While these two WRKYs were required for SA-induced *PR1* gene expression, they also did not control NHP-elicited potentiation of SA responses (Fig 6). Lastly, we examined the roles of WRKY38 and WRKY62, the expression of which are induced by NHP (*33*). Crucially, these two WRKYs are not required for SA-induced resistance (Fig S4), but are essential for activation of true SAR induced by an avirulent pathogen (*14*). Indeed, we discovered that WRKY38/62 are critical for NHP-elicited priming of the SA-responsive transcriptome and for amplifying SA-induced immunity (Figs 6 and 7).

WRKY38 and WRKY62 are type III WRKY family transcription factors with similar domain structures (*40*). Interestingly, they have both been shown to physically interact with the histone deacetylase, HDA19 (also known as HD1)(*40*). HDA19 expression is induced in response to SA, jasmonic acid and pathogen infection in an NPR1-independent manner (*40*), and its expression levels are anticorrelated with histone acetylation levels (*41*). Genetic analysis suggests that HDA19 plays important roles in regulating immune gene expression, including by repressing SA biosynthesis and downstream gene expression (*40-42*) Thus, it is plausible that during SAR, NHP-induced WRKY38/62 utilise the epigenetic modifier activity of HDA19 to prime the SA-responsive transcriptome in systemic tissues. Indeed, epigenetic histone modifications, including acetylation and methylation, have previously been linked to immune priming. Chemical or pathogen-induced SAR led to H3 and H4 acetylation as well as NPR1-dependent methylation at immune genes in systemic tissues (*43*). Moreover, the Jumonji (JMJ) domain-containing H3K4 demethylase, JMJ14, was found to regulate NHP biosynthesis and the pathogen-induced expression of immune genes (*44*). It has also been shown that transgenerational inheritance of priming occurs through wholescale changes in global DNA methylation (*45, 46*). Collectively, these findings suggests that gene priming involves epigenetic modifications that ensure rapid and amplified immune gene expression upon pathogen challenge. It will be interesting to assess in future to what extend WRKY38/62 affect histone modifications at NHP-primed, SA-responsive genes.

In summary, our findings represent a leap forward in understanding the mechanisms by which mobile immune signals establish SAR. We demonstrated that the major roles of AzA and NHP are to extensively modulate and potentiate the SA-responsive transcriptome, tailoring it to the needs of primed systemic tissues. Our study could conceivably function as a blueprint for studying the roles of other mobile immune signals (*e*.*g*. free radicals (*20*), glycerol-3-phosphate (*21*), dehydroabietinal (*22*)) in cross talking with immune hormones to establish durable systemic immunity. By identifying WRKY38/62 as novel regulators of NHP-mediated priming of SA responses and SA-induced immunity, we unveil the intriguing possibility that the wider WRKY transcription factor family may be involved in the activation of systemic immunity by other mobile immune signals.

## Materials and Methods

### Plant growth conditions and chemical treatments

*Arabidopsis thaliana* plants were of the Columbia-0 (Col-0) ecotype. Seeds were sterilised by washing with 96% ethanol, followed by a 15-min incubation in 50% bleach supplemented with 0.1% Triton. Seeds were then washed 5 times with sterilised water and stratified for 2-4 days in darkness at 4°C. For seedling experiments, seeds were plated on Murashige and Skoog agar (*47*) and grown at 22°C under long-day conditions (16h photoperiod; 06:00am to 22:00pm) with lighting intensity of 70-100 μmol m^-2^sec^-1^ for 10-14 days. For experiments on adult plants (28 days old), seeds were germinated at high humidity on an autoclaved soil mix of peat moss, vermiculite and sand at a 4:1:1 ratio. Subsequently, 10-12 day-old seedlings were transplanted to larger pots and grown on. Mutant genotypes used in this study are *35S:NPR1-GFP* (in *npr1-1*)(*48*), *npr3 npr4* (*6*), *cul3a/b (14), ube4* (SAIL_713_A12)(*15*) and *wrky38 wrky62* (SAIL_749_B02 and SM_3_38820)(*14*).

For AzA treatments, seedlings were immersed in 10 ml of 5 mM MES (pH5.6; Acros Organics #327765000) supplemented without (mock) or with 1 mM AzA (Azelaic acid 98%, Acros Organics #401521000) for 18h (16:00pm to 10:00am), then moved into 10 ml of 0.5 mM SA (sodium salicylate ≥99.5%, Sigma-Aldrich #S3007) for 6h (10:00am to 16:00pm). For NHP treatments, seedlings were treated by immersion in 10 ml of 1 mM NHP (Generon #HY-N7379) for 18h (16:00pm to 10:00am), then moved into 10 ml of 0.5 mM SA for 6h (10:00am to 16:00pm). For CHX assays plants were treated with NHP and SA as above, but 2h after addition of SA, 100µM CHX (Sigma-Aldrich#C7698) was added. Samples were taken immediately after adding CHX and thereafter at 2h, 4h and 6h timepoints.

### RNA extraction and qPCR

For RNA extractions, seedlings were grown for 10-14 days on MS plates and treated as described above. To extract RNA, seedlings were frozen in liquid nitrogen and ground into a fine powder. Equal volumes of phenol:chloroform:isoamlyalcohol (25:24:1) and an RNA extraction buffer (100 mM LiCl, 100 mM Tris (pH 8), 10 mM EDTA, 1% SDS) were added and the samples vortexed. The samples were then centrifuged at 13,300 rpm at 4°C for 5mins in a benchtop centrifuge before the aqueous phase was added to an equal volume of chloroform:isoamylalcohol (24:1) and briefly vortexed. This step was repeated and the aqueous phase incubated overnight with 1/3 volume 8M LiCl at 4°C. The precipitate was pelleted by centrifugation at 13,300 rpm at 4°C for 15 mins and the pellet washed with ice-cold 70% ethanol before rehydrating by resuspension in 400 µl H_2_0. The RNA was then precipitated by adding 1 ml ice-cold 96% ethanol and 40µl 3M NaAc (pH 5.2), and incubating for 1h at -20°C. Finally, purified RNA was pelleted by centrifugation at 13,300 rpm at 4°C for 15 mins, washed with ice-cold 70% ethanol, briefly air-dried, and resuspended in 25µl sterilised water.

RNA concentrations were quantified using a NanoDrop (ThermoFisher #ND-LITE-PR) and cDNA was synthesised using SuperScript II reverse transcriptase (ThermoFisher #18064022), according to the manufacturer’s instructions. cDNA was diluted 20-fold before qPCR analysis in which 4 µl cDNA, 5 µl SYBR green master mix (ThermoFisher #4309155) and 0.5 µl of each 10 µM primer (Table S4) were mixed. The samples were run in a qPCR thermocycler (ThermoFisher #4376600) with the following conditions: 95°C for 10min, (95°C for 15s, 60°C for 1min) x 40. The data were analysed using the ΔΔCt method (*49*).

### Disease assays

For disease assays, adult plants were grown on soil until 4 weeks old. They were then treated by spraying with 1mM AzA or 1 mM DL pipecolic acid (99%, Acros Organics #131280250) 48h before infection, followed by 0.5 mM or 0.05 SA 24 hours before infection. At 72h before infection, *Pseudomonas syringae* pv. *maliculicola* (*Psm*) ES4326 stocks stored at -80°C were streaked onto LB agar containing 10 mM MgCl_2_ and streptomycin (100 µg/ml). From a single colony, a liquid culture of LB containing 10 mM MgCl_2_ and streptomycin (100 µg/ml) was grown overnight. Bacterial cells were collected by centrifugation and resuspended in 10 mM MgCl_2_ to 5 x 10^6^ colony forming units/ml. Bacteria were pressure-infiltrated into leaves with a maximum of 2 infiltrated leaves per plant. At 3 or 4 days after infection, 8 representative leaves per sample were photographed before circular leaf discs (5 mm diameter) were cut out and ground fresh in 10 mM MgCl_2_. The resulting suspensions were serial diluted and streaked on LB agar plates containing 10 mM MgCl_2_ and streptomycin (100 µg/ml). Plates were incubated at room temperature and colony-forming units counted after 2-3 days. Three biological repeats of each assay were combined for analyses and graphing.

### Protein extraction and Western blotting

For protein experiments, seedlings were grown for 10-14 days on MS plates and treated as described above. To extract total protein, seedlings were frozen in liquid nitrogen and ground into a fine powder. Next, 2X (w/v) protein extraction buffer (50 mM Tris-HCl (pH 7.5), 150 mM NaCl, 5 mM EDTA, 0.1% Triton X-100, 0.2% Nonidet P-40, 1x protease inhibitor cocktail (142 nM TPCK, 135.5 nM TLCK, 0.5 nM PMSF)) was added and samples vortexed. Cell debris was pelleted by centrifugation at 13,300 rpm at 4°C for 15 mins, after which the supernatant was diluted into 1X protein sample buffer (40% glycerol, 240 mM Tris(pH 6.8), 8% SDS, 0.04% Bromophenol blue) supplemented with 50 mM DTT (Melford #D11000) and heated at 70°C for 10 mins. Samples were run on a 10% SDS-PAGE gel and transferred onto nitrocellulose membrane at 20V overnight. Membranes were blocked in 1X phosphate-buffered saline (PBS) with 5% non-fat milk powder and 0.1% TWEEN (SLS #che3852). Proteins were detected using α-GFP (1:5000 dilution, Roche #11814460001), α-S2 (1:2000 dilution, Abcam #98865) and α-RPN10 (1:2000 dilution, Abcam#56851) primary antibodies, followed by either anti-mouse (1:2000 dilution, Cell Signalling #7076) or anti-rabbit (1:2000 dilution, Cell Signalling #7074) secondary antibodies. Blots were visualised on a LICOR Odyssey Fc imaging system, using SuperSignal West Pico PLUS chemiluminescent substrate (ThermoFisher #34580).

### RNA sequencing analysis

For RNA-seq analysis, seedlings were grown and treated with AzA or NHP followed by treatment with SA as described above, then total RNA was extracted as described above. RNA integrity was confirmed by Bioanalyzer (Agilent 2100) and treatment efficacy was confirmed by qPCR for reporter genes as described above. High quality samples were then submitted to BGI Genomics (Hong Kong) for sequencing. Transcripts were quantified from raw reads using Kallisto following the authors instructions (*50*). Differentially expressed genes were identified from transcripts per million values using Sleuth following the authors instructions (*51*).

SA expressed genes were identified through linear modelling (Expression ∼ SA treatment, p ≤ 0.05), after which NHP primed behaviour was identified by hierarchical clustering with hclust and visualisation with ggplot2 (*52*), ggdendro (*53*) and ComplexHeatmap (*54*). Some clusters did not resolve as well as others, so were run through hclust again for better resolution. Cluster behaviour was scored through cutoffs when expression was scaled from 0 to 1, *i*.*e*. “NHP primed” genes were identified with “NW ≤ 0.2 and WS ≤ 0.8”. Clustering reliability was confirmed with heatmaps. When all groups had been assigned for the wild type and *wrky38/62*, Sankey diagrams were plotted with ggsankey (*55*). GO term plots were produced using the coriell package (*56*). Other R packages used during data processing, analysis and visualisation were tidyverse (*57*), factoextra (*58*), gridExtra (*59*), viridis (*60*), circlize (*61*), gridtext (*62*), cowplot (*63*), scales (*64*) and ggnewscale (*65*).

Scripts used for RNA-seq analysis and transcription factor associations are available at https://github.com/RobertOskar/Mason_2025. RNA-seq data have been deposited in Array Express at EMBL-EBI under accession code E-MTAB-14632 (AzA), E-MTAB-14631(NHP - WT) and E-MTAB-14847(NHP – *wrky38/62*).

### Statistical analysis

All statistical analyses were performed as described in the relevant figure legends. All qPCR graphs have error bars displaying ±SD. Disease assay significance was determined using Tukey post hoc analysis of variance (ANOVA) tests with α = 0.05. SA regulation in RNA-seq was determined using linear models (Expression ∼ SA treatment, α ≤ 0.05). Significant transcription factor association with primed gene categories was determined by comparing total number of genes targeted by each TF and the number of primed genes targeted by each TF using Chi-squared and Fishers exact test with α ≤ 0.05. RNA-seq, qPCR and protein quantification were performed on three replicas per sample and disease assays were performed on 24 separate individuals.

## Acknowledgements

We thank Prof. Jürgen Zeier for valuable discussions as well as Dr. Beatriz Orosa, Dr. Lucas Frungillo, and Dr. Hee-Kyung Ahn and their research teams for valuable feedback. We also thank Dr. Sophie Haupt and her team at the Plant Growth Facilities of the School of Biological Sciences, University of Edinburgh.

## Author contributions

Conceptualization: ROM, SHS

Methodology: ROM, SHS

Investigation: ROM, HG

Visualization: ROM

Supervision: SHS

Administration: HG, SHS

Writing: ROM, SHS

## Data and materials availability

All data are available in the main text or the Supplementary Materials except for RNA-seq data, which have been deposited in Array Express at EMBL-EBI under accession codes E-MTAB-14632, E-MTAB-14631 and E-MTAB-14847. Materials can be provided by S.H.S. pending scientific review and a completed material transfer agreement. Requests for the materials should be submitted to S.H.S.

