## Supplemental figures and tables for "Mobile immune signals potentiate salicylic acid-mediated plant immunity via WRKY38/62 transcription factors"

Robert O Mason *et al.*

**This PDF file includes:**

Figs. S1 to S4
Tables S1 to S4

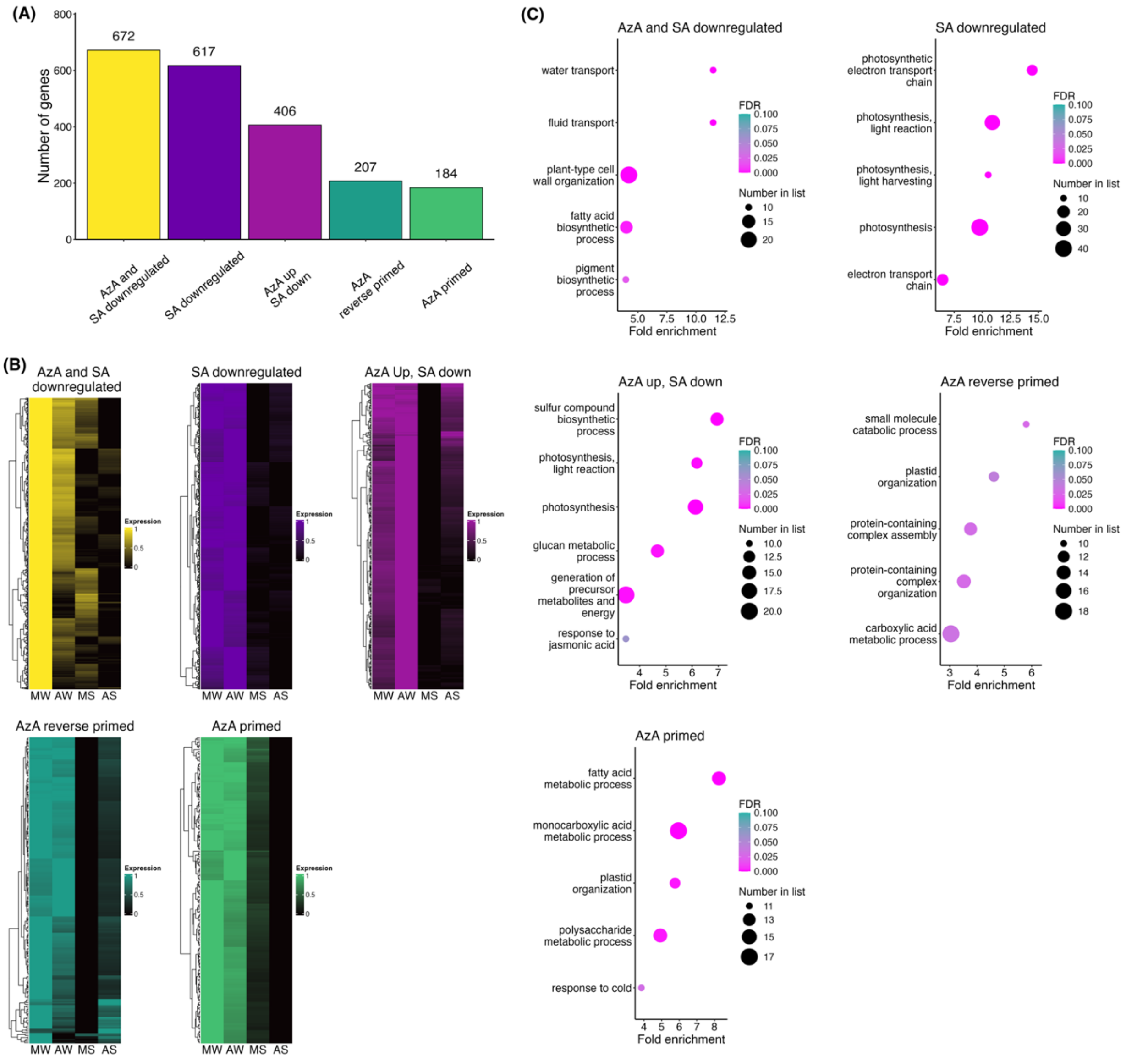


**Fig. S1. AzA differentially affects SA-repressed genes.** (**A**) RNA-seq clusters of AzA-modified, SA-repressed genes. Clusters were identified and assigned by hierarchical clustering using the hclust R package. (**B**) Heatmaps of clustered genes with expression patterns across the four treatment conditions (MW: mock + water, AW: AzA + water, MS: mock + SA, AS: AzA + SA). (**C**) Top 5 most enriched GO terms in each cluster ($p \leq 0.05$, FDR $\leq 0.1$). Fourteen-day-old wild-type seedlings were treated by immersion in 5 mM MES (pH 5.6) with or without 1 mM AzA for 18 hours, followed by treatment with water or 0.5 mM SA for 6 hours. Total RNA was extracted and the expression of SA-responsive genes measured by RNA seq.

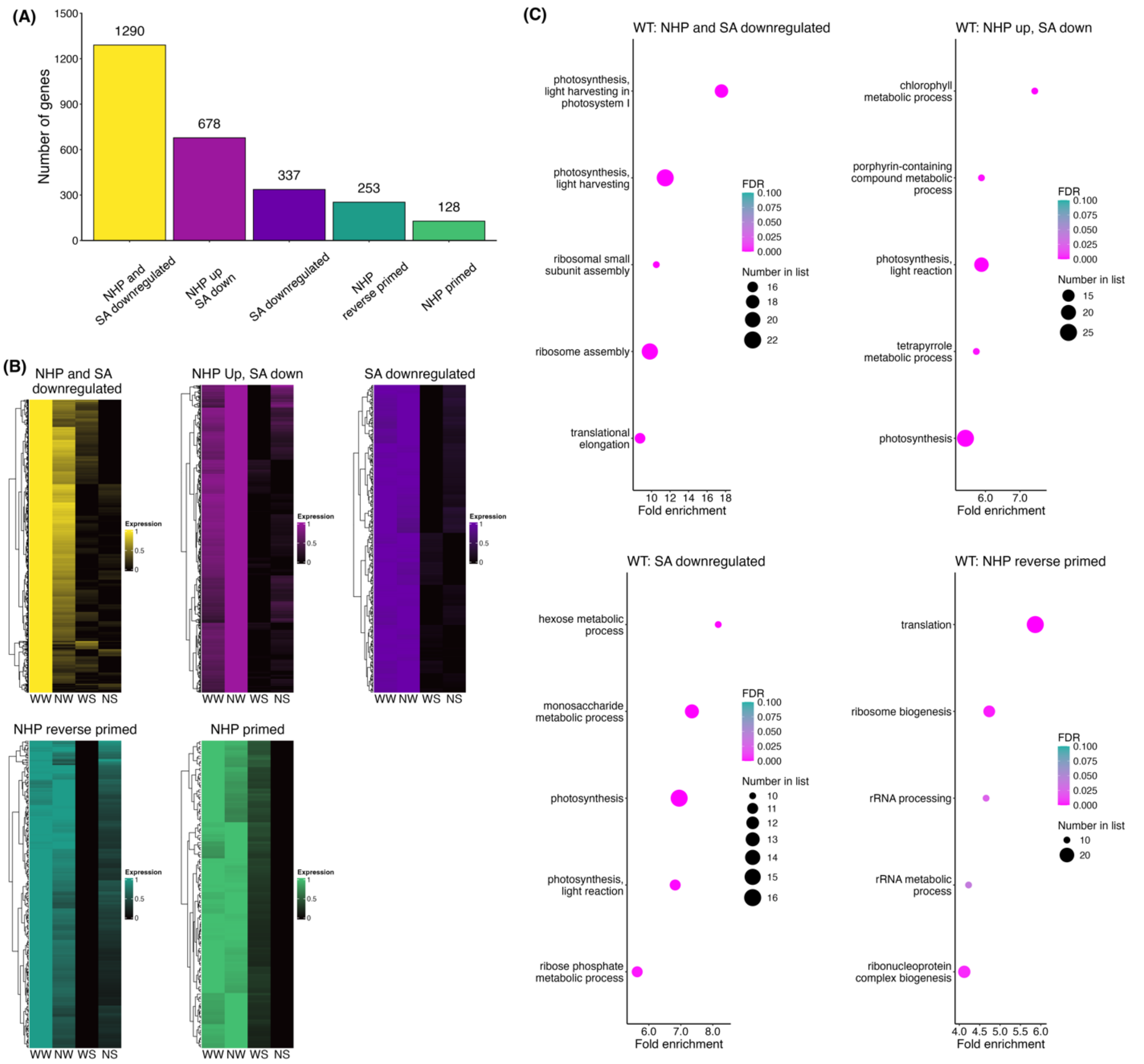


**Fig. S2. NHP differentially affects SA-repressed genes.** (A) RNA-seq clusters of NHP-modified, SA-repressed genes. Clusters were identified and assigned by hierarchical clustering using the hclust R package. (**B**) Heatmaps of clustered genes with expression patterns across the four treatment conditions (WW: water + water, NW: NHP + water, WS: water + SA, NS: NHP + SA). (**C**) Top 5 most enriched GO terms in each cluster ($p \leq 0.05$, FDR $\leq 0.1$). For all experiments, 14-day-old wild-type seedlings were treated by immersion in water or 1 mM NHP for 18 hours, followed by treatment with water or 0.5 mM SA for 6 hours. Total RNA was extracted and the expression of SA-responsive genes measured by RNA seq.

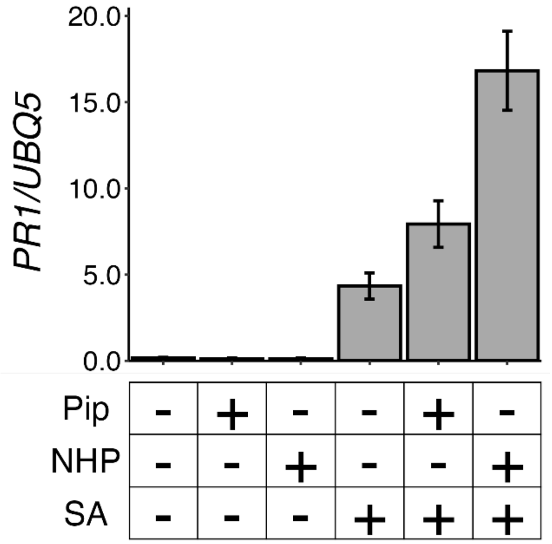


**Fig. S3. Both Pip and NHP prime SA-induced expression of PR1.** Fourteen-day-old wild-type seedlings were treated by immersion in water, 1mM Pip (*i.e.* NHP precursor) or 1 mM NHP for 18 hours, followed by treatment with water or 0.5 mM SA for 6 hours. Total RNA was extracted and the expression of SA-responsive genes measured by qPCR and normalised against constitutively expressed *UBQ5*.

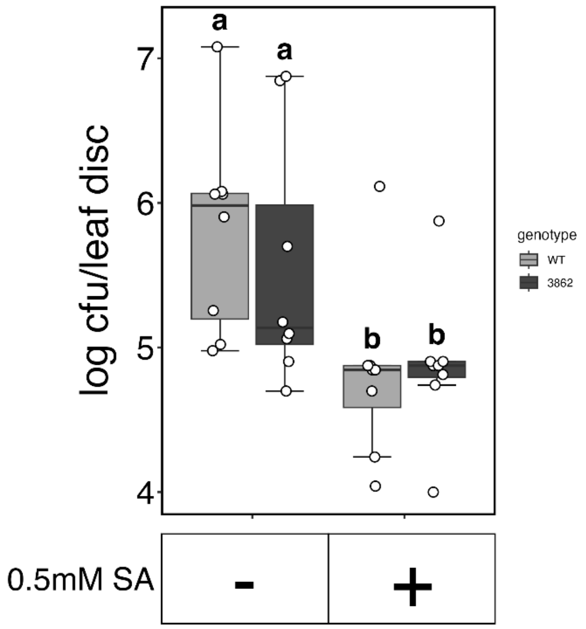


**Fig. S4. SA-induced disease resistance in *wrky38/62* mutants is comparable to wild type.** Twenty-eight day-old wild-type or *wrky38/wrky62* plants were sprayed with water or 0.5 mM SA, 24 hours before infection. Leaves were then infiltrated with 5 x $10^6$ colony forming units (cfu)/ml *Psm* ES4326. Leaf discs were analysed for bacterial growth two days post infection. Error bars represent interquartile range x 1.5 limits, while letters denote statistically significant differences between samples (Tukey ANOVA; α = 0.05, n = 8).

**Table S1. Differentially expressed transcripts identified in AzA priming RNA seq.** (**A**) Genes up- and downregulated by AzA alone. (**B**) SA up- and downregulated genes and their AzA priming response.

**Table S2. Differentially expressed transcripts identified in NHP priming RNA seq.** (**A**) SA up- and downregulated genes and their NHP priming response. (**B**) Genes up- and downregulated by NHP alone.

**Table S3. Transcription factor binding sites in NHP-primed genes.** Using published DAPseq datasets, transcription factor targets in all differentially expressed genes were compared to NHP primed targets. Significant enrichment was determined using Chi-squared and Fishers exact test ($\alpha \leq 0.05$).

**Table S4. All primers used in this study.**
